# GABA neurons in the sublaterodorsal tegmental (SLD) nucleus suppress wakefulness in wild-type and narcoleptic mice

**DOI:** 10.1101/2025.07.10.664050

**Authors:** HanHee Lee, Jimmy J. Fraigne, John H. Peever

## Abstract

The sleep-wake cycle is generated by competing neural circuits that control the oscillation between wakefulness, rapid eye movement (REM) sleep, and non-REM (NREM) sleep. While the sublaterodorsal tegmental nucleus (SLD) is recognized for its role in REM sleep generation, the functional contribution of its GABAergic neurons (SLD^GABA^) to sleep-wake regulation remains poorly understood. Here, we found that SLD^GABA^ neurons function as a suppressor of wakefulness in both wild-type and narcoleptic mice. Optogenetic silencing of SLD^GABA^ neurons rapidly induced robust wakefulness, while enhancing cortical and motor activity. Conversely, optogenetic activation of these neurons suppressed wakefulness and promoted NREM sleep. We found that SLD^GABA^ neurons project extensively to wake-promoting brain regions, providing an anatomical basis for their wake-suppressing effects. Importantly, we discovered that SLD^GABA^ neurons play a pathological role in narcolepsy: their activation in orexin-deficient mice triggered characteristic sleep attacks—rapid intrusions of NREM sleep during active wakefulness— while silencing these neurons rescued animals from both sleep attacks and cataplexy. Collectively, these findings establish SLD^GABA^ neurons as a key regulator of arousal state transitions and identify them as a novel therapeutic target for the treatment of narcolepsy.

## INTRODUCTION

Current models propose that sleep-wake regulation is orchestrated by distinct neuronal populations and circuits that compete for each arousal state control (i.e., wakefulness, NREM and REM sleep). The brainstem contains critical parts of this regulatory network, yet the precise mechanisms governing arousal state transitions remain incompletely understood^1–3^. Understanding these circuits is essential not only for basic sleep biology but also for developing treatments for sleep disorders.

The sublaterodorsal tegmental nucleus (SLD), located in the dorsal pons near key arousal centers, including the locus coeruleus and parabrachial nucleus, plays a strategic role within the sleep-wake regulatory network^4–6^. While the SLD is typically considered as the generator of REM sleep, emerging evidence suggests it also influences wakefulness. Pharmacological studies demonstrate that SLD activation promotes REM sleep while suppressing wakefulness, whereas its inhibition prevents REM sleep and enhances wakefulness^7,8^. This control over both REM sleep and wakefulness positions the SLD as a critical hub in arousal state regulation.

The SLD contains both glutamate and GABA neurons. Glutamate SLD (SLD^GLU^) neurons are well-established promoters of REM sleep and its associated muscle atonia^9–11^. In contrast, the functional role of GABA SLD neurons (SLD^GABA^) remains speculative. For example, some studies suggest that SLD^GABA^ neurons suppress wakefulness by inhibiting the ascending reticular activating syste ^2,12,13^, while other studies suggest that they regulate REM sleep^14^ or have minimal effect on sleep-wake states^11^. These conflicting findings highlight a critical gap in our understanding of the function of SLD^GABA^ neurons in sleep-wake regulation, and underscore the need for precise, cell-type-specific approaches (i.e., genetic tagging and optogenetic) to resolve their role in sleep-wake control..

The clinical relevance of understanding SLD function extends to narcolepsy, a debilitating neurological disorder characterized by pathological intrusions of sleep into wakefulness. Narcolepsy results from the loss of orexin (also called hypocretin) signalling, which normally stabilizes wakefulness. Narcolepsy is characterized by pathological disruptions of wakefulness, manifesting as sleep attacks—sudden intrusion of NREM sleep into wakefulness—and cataplexy—intrusions of REM sleep muscle atonia into wakefulness^9,15^. Neuroanatomical and pharmacological evidence indicates that orexin neurons extensively innervate and interact with the SLD, and the loss of orexin signalling profoundly impacts SLD function^9,14,16–18^. Our recent work demonstrated that SLD^GLU^ neurons, which promote REM sleep muscle atonia in healthy mice (i.e., orexin^+/+^), drive cataplexy in narcoleptic mice (i.e., orexin^-/-^) ^9,19^. This suggests that the SLD may be a central hub where orexin loss triggers the characteristic symptoms of narcolepsy.

Despite the clinical importance of the SLD in narcolepsy, the specific contribution of SLD^GABA^ neurons to the disorder’s pathophysiology remains unknown. Given their proposed role in suppressing wakefulness, we hypothesized that SLD^GABA^ neurons might contribute to the sleep attacks that affect narcoleptic patients. Understanding this relationship could reveal new therapeutic approaches for this disorder.

Here, we used optogenetics, electrophysiology, and genetically-assisted circuit mapping to establish the role of SLD^GABA^ neurons in sleep/wake control. First, our findings reveal that SLD^GABA^ neurons function as powerful suppressors of wakefulness likely through direct projections to wake-promoting brain regions. Then, we show that these neurons play a direct role in triggering narcoleptic sleep attacks, as their activation triggers pathological sleep intrusions while their silencing provides therapeutic rescue. Our findings not only resolve a longstanding question about the function of SLD^GABA^ neurons but also identify a potential intervention target to resolve the excessive sleepiness of narcolepsy.

## RESULTS

### Silencing SLD^GABA^ neurons promotes wakefulness

To test the role of SLD^GABA^ neurons in sleep-wake regulation, we used optogenetics to silence their activity across natural sleep-wake states in wild-type mice. We did this by bilaterally injecting either AAV-EF1a-DIO-eArch3.0-eYFP or AAV-EF1a-DIO-mCherry (control) into the SLD of male *VGAT*-Cre mice (**Figure 1A and 1B**). Then, we implanted optic fibers above either Arch-or mCherry-expressing SLD^GABA^ neurons for laser manipulation **(Figure S1A and S1B)**. EEG and EMG electrodes were implanted to monitor and identify each arousal state.

**Figure 1.**
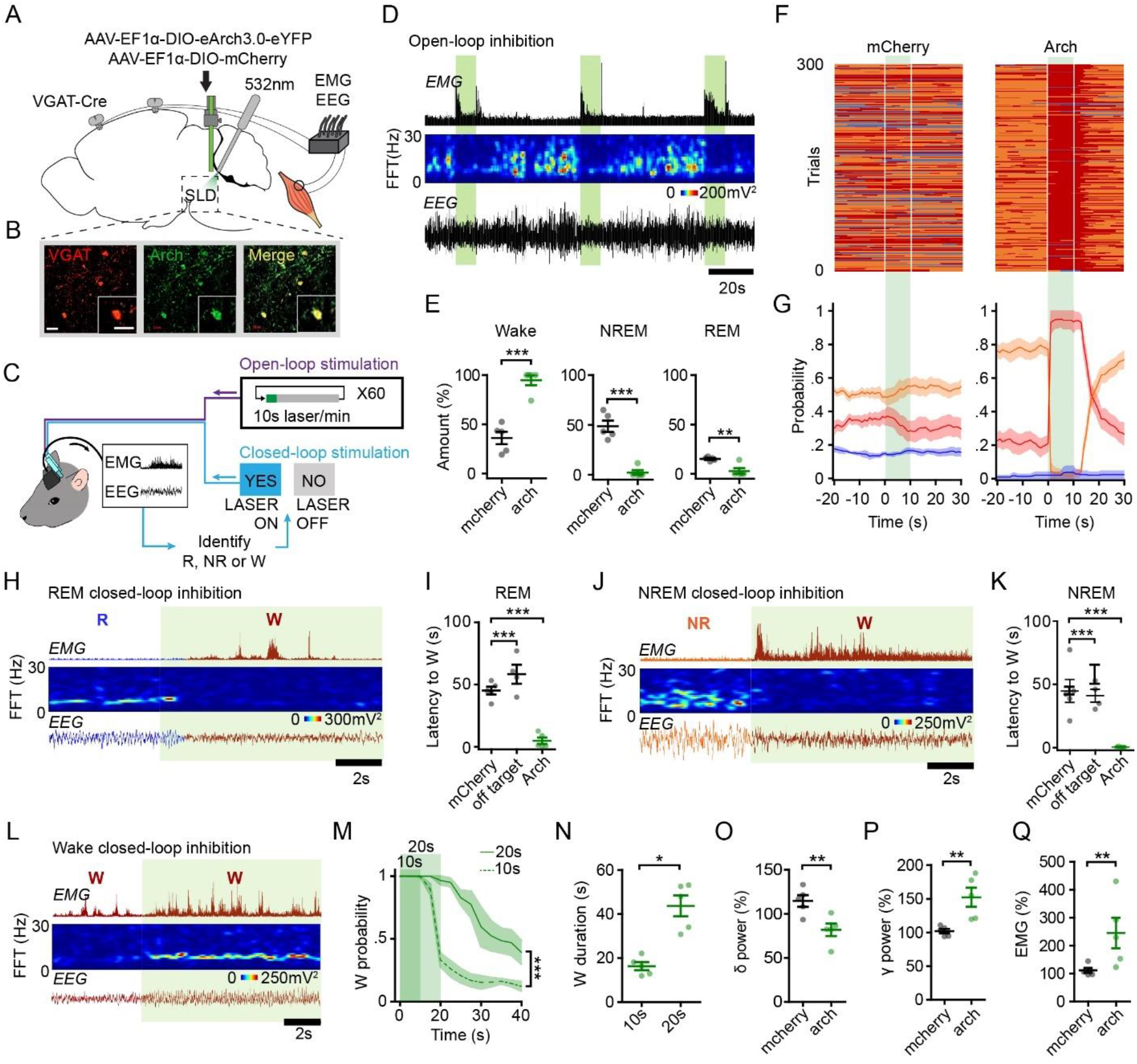
Silencing SLD^GABA^ neurons promotes wakefulness. **(A)** A schematic showing optogenetic silencing of SLD^GABA^ neurons coupled with EEG and EMG recordings. **(B)** Microscope images showing the expression of Arch (green) from *VGAT (*red) neurons in the SLD. **(C)** A schematic of open-loop and closed-loop stimulation paradigm. **(D)** Example polysomnogram recording with open-loop stimulation (i.e., 10s laser stimulation every 60s for 1 hour) of Arch-expressing SLD^GABA^ neurons. Shown are EMG amplitude, EEG spectrogram and EEG raw traces. **(E)** Mean percentages of wake, NREM, and REM sleep during 10s open-loop stimulation (mCherry n=5 and Arch n=5; unpaired t-test). **(F)** Distribution of sleep-wake states in all laser trials aligned by the time of laser onset at t = 0s (mCherry n=5 and Arch n=5). **(G)** Probability of wake, NREM, and REM sleep before, during, and after 10s open-loop stimulation (mCherry n=5 and Arch n=5). **(H)** Example polysomnogram recording with closed-loop stimulation of Arch-expressing SLD^GABA^ neurons during REM sleep. **(I)** Latency to wake from REM sleep upon laser stimulation (mCherry n=5, off-target n=4 and Arch n=5, one-way ANOVA with Tukey’s Multiple Comparison test). **(J)** Example polysomnogram recording with closed-loop stimulation of Arch-expressing SLD^GABA^ neurons during NREM sleep. **(K)** Latency to wake from NREM sleep upon laser stimulation (mCherry n=5, off-target n=4 and Arch n=5, one-way ANOVA with Tukey’s Multiple Comparison test). **(L)** Example polysomnogram recording with closed-loop stimulation of Arch-expressing SLD^GABA^ neurons during wakefulness. **(M)** Probability of wakefulness in response to 10s and 20s laser stimulation of Arch-expressing SLD^GABA^ neurons (n=5, two-way ANOVA with Bonferroni post-test). **(N)** Duration of wakefulness in response to 10s and 20s laser stimulation of Arch-expressing SLD^GABA^ neurons (n=5, paired t-test). **(O-P)** Mean δ and γ EEG power during closed-loop stimulation for wakefulness (mCherry n=5, Arch n=5, unpaired t-test). **(Q)** Mean EMG activity during closed-loop stimulation for wakefulness (mCherry n=5 and Arch n=5; unpaired t-test). Brain wave bands: δ (delta, 0.5-4Hz) and γ (gamma, 30-100Hz). Green patches indicate time of laser stimulation. All error bars and shades represent ±s.e.m. *\* p<0.05, ** p<0.01, *** p<0.001 indicate significant differences.* Scale bar 25um.

We first silenced the activity of SLD^GABA^ neurons using an open-loop protocol. We delivered light (532nm) continuously for 10s every minute for one hour then quantified the changes in arousal states **(Figure 1C)**^20^. We found that the silencing of Arch-expressing SLD^GABA^ neurons reliably induced episodes of wakefulness (**Figure 1D**). To quantify this effect, we aligned the laser trials from all mice by the time of laser onset and then calculated the amount and probability of each arousal state across time (**Figure 1F**). We found that optical silencing of Arch-expressing SLD^GABA^ neurons induced a robust increase in the amount and probability of wakefulness and a complementary decrease in both NREM and REM sleep (**Figure 1E-G**). Laser manipulation of mCherry-expressing SLD^GABA^ neurons (control) had no effect on the arousal states (**Figure 1E-G, Figure S2)** nor did light applied to regions adjacent to Arch-expressing SLD^GABA^ neurons (i.e., off-target optic fibers) (**Figure 1I and 1K**). This finding supports that silencing of SLD^GABA^ neurons is inducing wakefulness rather than the laser itself.

To further determine the role of SLD^GABA^ neurons, we applied a closed-loop protocol whereby we selectively silenced SLD^GABA^ neurons during each arousal state. To do this, we monitored real-time EEG/EMG activity and silenced these neurons as soon as spontaneous episodes of either wakefulness, NREM, or REM sleep occurred **(Figure 1C)**. We found that silencing SLD^GABA^ neurons either during NREM or REM sleep induced instantaneous transitions into alert and motorically engaged wakefulness (**Figure 1H-K & Figure S1C-F & Video S1**). We also found that silencing SLD^GABA^ neurons during wakefulness sustained wakefulness. A prolonged silencing of SLD^GABA^ neurons (20s vs. 10s) induced a longer-lasting wakefulness and maintained a higher probability of wakefulness over time (**Figure 1L-N**). These findings demonstrate that silencing SLD^GABA^ neurons not only initiates wakefulness from sleep but also sustains wakefulness.

Next, we investigated whether silencing SLD^GABA^ neurons not only triggers wakefulness but also can enhance the features of wakefulness. To do this, we silenced SLD^GABA^ neurons while the animals were already awake. We found that silencing SLD^GABA^ neurons during wakefulness increased the power of wake-related cortical activity (i.e., gamma, 30-100Hz; **Figure 1P**) while simultaneously decreased sleep-related cortical activity (i.e., delta, 0.5-4Hz; **Figure 1O & Figure S1H-L**). In addition, we found that silencing SLD^GABA^ neurons during wakefulness also enhanced muscle activity (i.e., increased EMG activity) (**Fig 1Q & Figure S1G**). Together, our findings show that silencing SLD^GABA^ neurons not only triggers wakefulness but also enhances brain and muscle activity that characterize active arousal. These findings suggest that SLD^GABA^ neurons play a powerful role of suppressing wakefulness and its wake-related features.

### Activation of SLD^GABA^ neurons suppresses wakefulness

To further examine the role of SLD^GABA^ neurons in sleep-wake control, we optically activated these neurons and assessed their impact on each arousal state (**Figure 2**). To do this, we bilaterally injected AAV-EF1a-DIO-ChETA-eYFP or AAV-EF1a-DIO-mCherry (control) into the SLD of male *VGAT*-*Cre* mice **(Figure 2A).** Optic fibers were positioned above SLD^GABA^ neurons and EEG/EMG readout was used to assess how the laser stimulation affects arousal states. Laser stimulation (478nm) was delivered in pulses (5ms) at 40Hz to match the natural firing rates of SLD^GABA^ neurons reported in the literature^21–23^.

**Figure 2.**
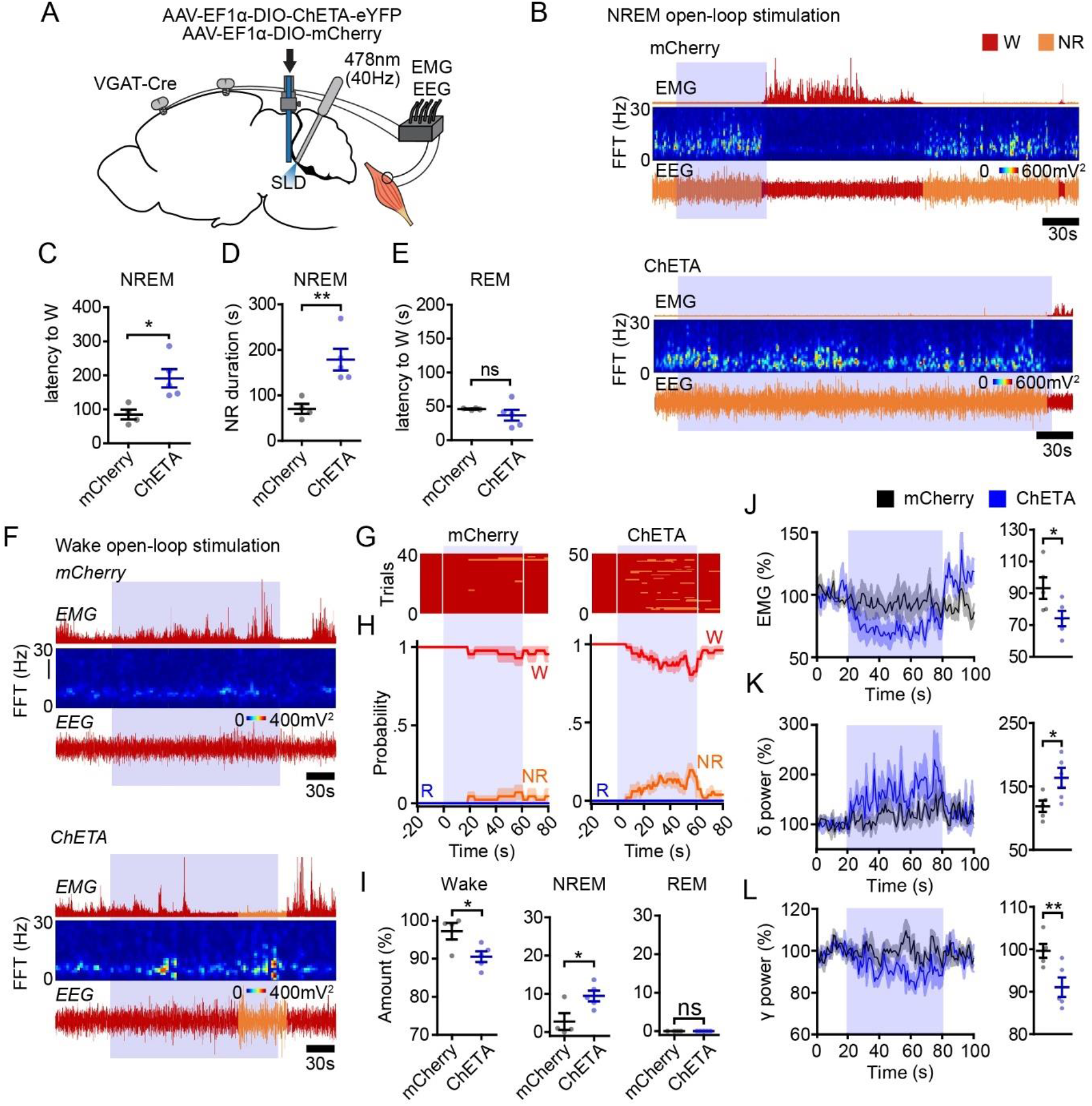
Activating SLD^GABA^ neurons suppresses wakefulness. **(A)** A schematic showing optogenetic activation (478nm, 40Hz, 5ms pulses) of SLD^GABA^ neurons coupled with EEG and EMG recordings. **(B)** Example polysomnograms with closed-loop stimulation of mCherry (top) and ChETA (bottom) expressing SLD^GABA^ neurons during NREM sleep. Shown are EMG amplitude, EEG spectrogram, and EEG raw traces. **(C)** Latency to wake from NREM sleep upon laser stimulation (mCherry n=4 and ChETA n=5; unpaired t-test). **(D)** Duration of NREM sleep upon laser stimulation (mCherry n=4 and ChETA n=5; unpaired t-tests). **(E)** Latency to wake from REM sleep upon laser stimulation (mCherry n=4 and ChETA n=5, unpaired t-test). **(F)** Example polysomnograms with closed-loop stimulation (60s) of mCherry (top) and ChETA (bottom) expressing SLD^GABA^ neurons during wakefulness. **(G)** Distribution of sleep-wake states in all laser trials aligned by the time of laser onset t = 0s (mCherry n=4 and ChETA n=5). Laser trials (60s) were collected from closed-loop stimulation for wakefulness. **(H)** Probability of wake, NREM, and REM sleep before, during, and after the 60s laser stimulation (mCherry n=4 and ChETA n=5**). (I)** Mean percentages of wake, NREM, and REM sleep during the 60s laser stimulation (mCherry n=4 and ChETA n=5; unpaired t-test) **(J) *LEFT***: EMG activity before, during, and after 60s laser stimulation from wakefulness (mCherry n=4 and ChETA n=5). ***RIGHT***: Mean EMG activity during the 60s laser stimulation (mCherry n=4 and ChETA n=5; unpaired t-test). **(K-L) *LEFT*:** δ and γ EEG activity before, during, and after 60s laser stimulation from wakefulness (mCherry n=4 and ChETA n=5). ***RIGHT*:** Mean δ and γ EEG activity during the 60s laser stimulation (mCherry n=4 and ChETA n=5; unpaired t-test). EEG bands: δ (delta, 0.5-4Hz) and γ (gamma, 30-100Hz). Blue patches indicate time of laser stimulations. All error bars and shades represent ±s.e.m. *\* p<0.05, ** p<0.01, *** p<0.001 indicate significant differences.*

Because silencing SLD^GABA^ neurons promotes wakefulness (**Figure 1**), we predicted that their activation would suppress wakefulness and induce sleep. Using a closed-loop protocol, we found that 40Hz laser stimulation of ChETA-expressing SLD^GABA^ neurons during NREM sleep significantly lengthened NREM sleep by preventing the entrance into wakefulness (**Figure 2B-D**). In contrast, we found that 40Hz laser stimulation during REM sleep did not prolong REM sleep (**Figure 2E)**. We also found that neither cortical nor motor activity was affected by 40Hz laser stimulation during NREM or REM sleep **(Figure S3B-E),** indicating that the effect of activating SLD^GABA^ neurons specifically prolongs the state of NREM sleep. Laser stimulation of mCherry-expressing SLD^GABA^ neurons had no effect on NREM or REM sleep (**Figure S2**).

Next, we found that 40Hz activation of SLD^GABA^ neurons suppressed the probability and amount of wakefulness by increasing NREM sleep but not REM sleep **(Figure 2 F-I)**. In addition, we found that activating SLD^GABA^ neurons reduced the cortical and motor indices of wakefulness. Specifically, we found that activation of SLD^GABA^ neurons during wakefulness significantly increased and decreased EEG δ and γ powers, respectively, while simultaneously decreasing levels of motor activity (i.e., EMG) **(Figure 2J-L & Figure S3 F-K)**. Together, these data show that the SLD^GABA^ neurons suppress wakefulness and the cortical and motor features that support it.

### SLD^GABA^ neurons project to wake-promoting brain areas

Having shown that SLD^GABA^ neurons suppress wakefulness, we next wanted to identify the projection patterns of SLD^GABA^ neurons. To do this, we injected a Cre-dependent anterograde viral tracer (AAV-EF1a-DIO-eArch3.0-eYFP) into the SLD of *VGAT*-*Cre* mice and then mapped the location of eYFP-expressing SLD^GABA^ axonal projections throughout the brain. We found that SLD^GABA^ axons predominantly project to brain regions that contain neurons associated with generating wakefulness and engaging cortical and motor activity **(Figure 3A)**^24–29^. Specifically, we identified eYFP-expressing axon projections in the medial septum, basal forebrain, lateral hypothalamus, tuberomammillary nucleus, supramammilary nucleus, interpeduncular nucleus, and the ventrolateral periaqueductal grey many of which play key roles in promoting wakefulness and/or inducing cortical activation **(Figure 3B-H)**. We also identified SLD^GABA^ axon projections in the vestibular nucleus and in the ventral horn of the spinal cord, which both play key roles in engaging motor activity **(Figure 3I and 3J)**^30–32^. These findings indicate that SLD^GABA^ neurons are anatomically positioned to modulate wakefulness, cortical activity, and motor activity, which is consistent with our optogenetic data.

**Figure 3.**
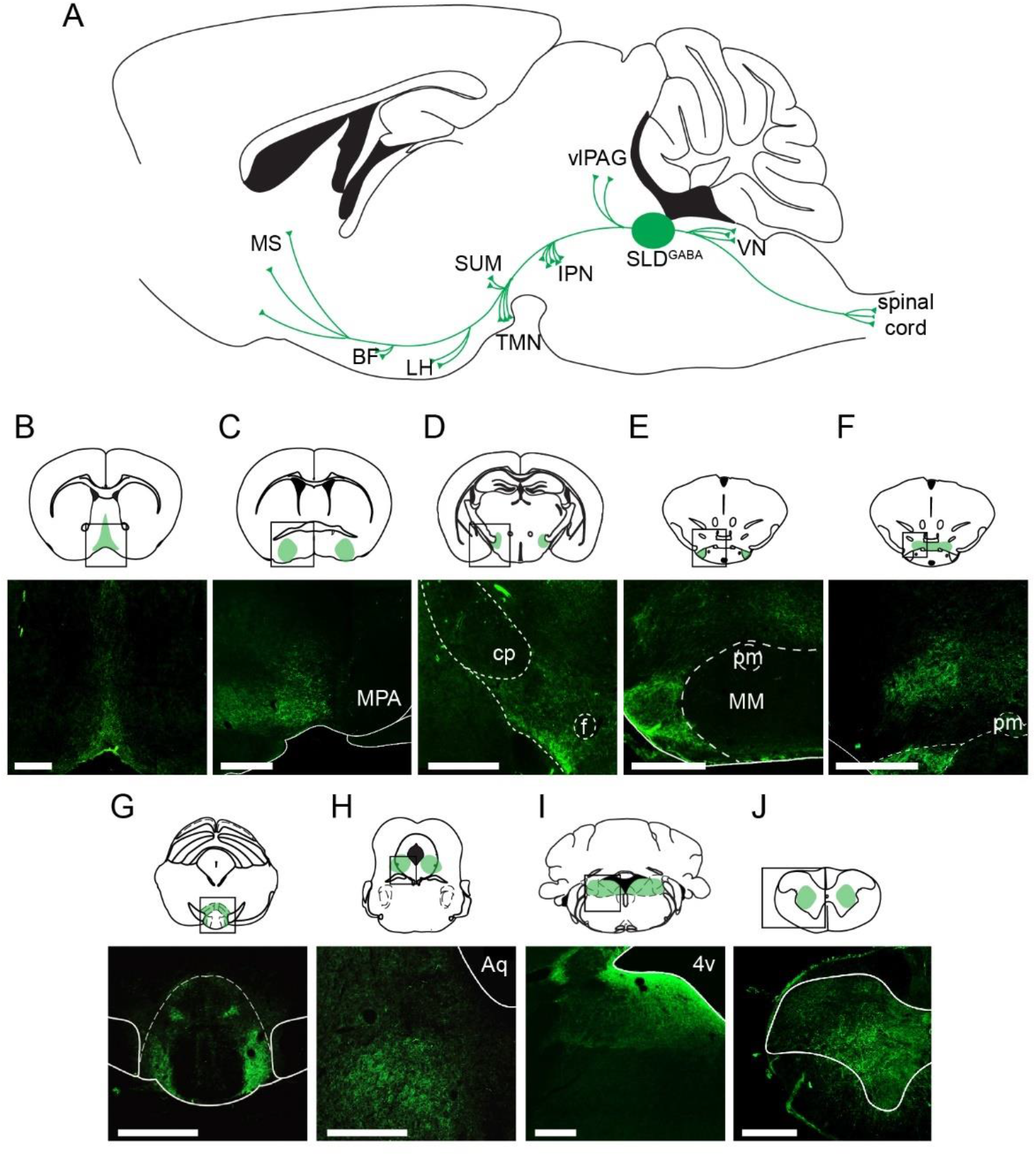
SLD^GABA^ neurons send axon projections to wake-promoting nuclei. **(A)** A sagittal schematic of the brain illustrating the axon distribution of SLD^GABA^ neurons. **(B-J)** Schematics (***TOP***) and microscope images (***BOTTOM***) showing coronal section of each brain nuclei that receives axon input from SLD^GABA^ neurons (green fluorescence). Abbreviation: medial septum (MS), basal forebrain (BF), medial preoptic area (MPA), lateral hypothalamus (LH), cerebral peduncle (cp), fornix (f), tuberomammillary nucleus (TMN), principle mamillary tract (pm), medial mammillary body (MM), supramammilary nucleus (SuM), interpeduncular nucleus (IPN), ventrolateral periaqueductal grey (vlPAG), aqueduct (Aq), sublaterodorsal tegmental nucleus (SLD), trigeminal motor nucleus (Mo5), vestibular nucleus (VN) and 4^th^ ventricle (4V). Scale bar 400um.

### Activation of SLD^GABA^ neurons triggers sleep attacks in *orexin*^-/-^mice

Orexin is a neurotransmitter produced by neurons in the lateral hypothalamus, which send axonal projections across the brain to support wakefulness^16^. In narcolepsy, the loss of orexin signals destabilizes wakefulness, making this state vulnerable to sleep disturbances^15,33–35^. Previous work demonstrate that loss of orexin impacts how SLD neurons control arousal states^14,16,17^. For example, our recent work demonstrated that glutamate neurons in the SLD, which generate REM sleep muscle atonia, promote cataplexy in narcoleptic mice. In this current study, we decided to investigate how the loss of orexin impacts the function of SLD^GABA^ neurons. We hypothesized that activation of the SLD^GABA^ neurons in narcoleptic mice will potently suppress wakefulness and induce sleep attacks. To test this hypothesis, we bilaterally injected AAV-EF1a-DIO-ChETA-eYFP or AAV-EF1a-DIO-mCherry into the SLD of male *VGAT*-Cre::*orexin*^-/-^ mice. This newly developed mouse line behaves identically to ‘pure’ *orexin*^-/-^ mice and exhibits both sleep attacks and cataplexy^36^. EEG/EMG electrodes and optic fibers were implanted to manipulate GABA neurons in the SLD while monitoring sleep-wake states in *VGAT*-Cre::*orexin*^-/-^ mice (**Figure 4A and 4B)**.

**Figure 4.**
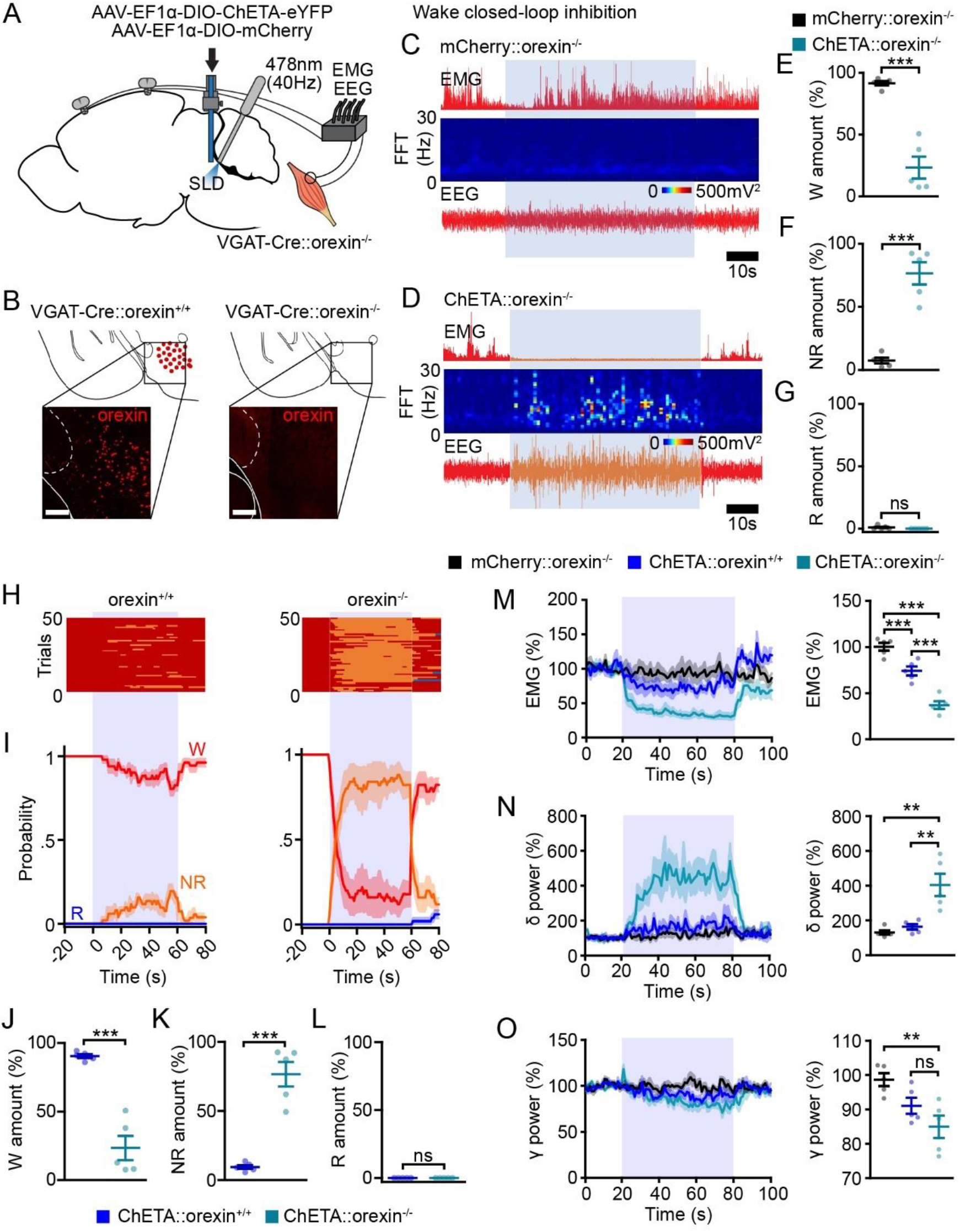
Activating SLD^GABA^ neurons in orexin^-/-^ mice potently suppresses wakefulness. **(A)** A schematic of optogenetic activation of SLD^GABA^ neurons coupled with EEG and EMG recordings in orexin^-/-^ mice. **(B)** Schematic and microscope images showing the absence of orexin neuropeptide (red) from the lateral hypothalamus (LH) in orexin^-/-^ (i.e., narcoleptic) mice. **(C-D)** Example polysomnogram with closed-loop stimulation (60s) of mCherry-expressing and ChETA-expressing SLD^GABA^ neurons during wakefulness. Shown are EMG amplitude, EEG spectrogram, and EEG raw traces. **(E-G)** Mean percentages of wake, NREM, and REM sleep during 60s closed-loop stimulation from wakefulness (mCherry::orexin^-/-^ n=4 and ChETA::orexin^-/-^ n=5; unpaired t-test). **(H)** Distribution of sleep-wake states in all laser trials aligned by the time of laser onset t = 0s (orexin^+/+^ n=5 and orexin^-/-^ n=5). Laser trials (60s) were collected from closed-loop stimulation for wakefulness. **(I)** Probability of wake, NREM and REM sleep before, during and after the 60s laser stimulation (orexin^+/+^ n=5 and orexin^-/-^ n=5). **(J-L)** Mean percentages of wake, NREM and REM sleep during 60s closed-loop stimulation from wakefulness (ChETA::orexin^+/+^ n=5 and ChETA::orexin^-/-^ n=5; unpaired t-test). **(M) *LEFT*:** EMG activity before, during, and after 60s closed-loop stimulation from wakefulness (mCherry::orexin^-/-^ n=5, ChETA::orexin^+/+^ n=5, and ChETA::orexin^-/-^ n=5). ***RIGHT*:** Mean EMG activity during the 60s laser stimulation (mCherry::orexin^+/+^ n=5, ChETA::orexin^-/-^ n=5, and ChETA::orexin^+/+^ n=5; unpaired t-test). **(N-O) *LEFT*:** δ and γ EEG activity before, during and after 60s closed-loop stimulation from wakefulness (mCherry::orexin^-/-^ n=5, ChETA::orexin^+/+^ n=5, and ChETA::orexin^-/-^ n=5). ***Right*:** Mean δ and γ EEG activity during the 60s laser stimulation (mCherry::orexin^-/-^ n=5, ChETA::orexin^+/+^ n=5, and ChETA::orexin^-/-^ n=5; unpaired t-test). EEG bands: δ (delta, 0.5-4Hz) and γ (gamma, 30-100Hz). Blue patches indicate time of laser stimulation. All error bars and shades represent ±s.e.m. *\*\* p<0.01 *** p<0.001 indicate significant differences.*

Using a closed-loop protocol, we optically activated SLD^GABA^ neurons (478nm, 5ms pulses at 40Hz) during each arousal state in orexin^-/-^ mice. We found that activation of SLD^GABA^ neurons during wakefulness triggered a sudden entrance into NREM sleep, which was not observed in the mCherry control group (**Figure 4C-G**). Intriguingly, we found that the effect of SLD^GABA^ neuron activation was different between *orexin*^-/-^ (i.e., narcoleptic) mice and *orexin*^+/+^ (i.e., wild-type) mice. Specifically, we show that the activation of SLD^GABA^ neurons during wakefulness had a stronger wake-suppressing effect in *orexin*^-/-^ than in *orexin*^+/+^ mice**(Figure 4H-L)**. In addition, we also found that activation caused a greater reduction in the cortical and motor activity in *orexin*^-/-^ mice **(Figure 4M-O & Figure S4O-T)**. Together, these findings demonstrate that SLD^GABA^ neuron activation suppresses wakefulness and the cortical and motor features that support it, but the loss of orexin signaling potentiates these effects, causing the typical sleep-attack symptoms observed in narcoleptic patients.

We found that the wake→NREM sleep transitions induced by the activation of SLD^GABA^ neurons in *orexin*^-/-^ animals was distinguishable from the wake→NREM sleep transitions observed in *orexin*^+/+^ animals by how quickly they occurred. In *orexin*^-/-^ mice, the activation consistently and rapidly terminated (i.e., ° 5 seconds) wakefulness into NREM sleep **(Figure 4H-L)**. The *orexin*^-/-^ mice remained in NREM sleep throughout the duration of activation (60s) and consistently and rapidly returned to wakefulness at the offset of activation **(Figure 4D, 4H and F4I)**, similar to what happened during naturally occurring sleep attacks. In addition, we found that activation in *orexin*^-/-^ mice could trigger a sleep attack regardless of the animal’s behaviour and location at the onset of activation. In *orexin*^+/+^ mice, wake→NREM sleep transitions occur when animals are settled in their nests^37^. However, in orexin^-/-^ mice, the activation of SLD^GABA^ neurons induced a rapid entrance into NREM sleep even when the animals were active outside of the nest (**Video S2**). **Video S2** is a typical behavioural example showing that activation triggers an abrupt and direct wake→NREM sleep transition during a period of active wakefulness. Abrupt intrusions of NREM sleep into wakefulness is the defining feature of sleep attacks in human and murine narcolepsy^33,37,38^; therefore, we reasoned that the activation of SLD^GABA^ neurons induced sleep attacks in *orexin*^-/-^ mice.

Next, we investigated whether the cortical activity during the activation-induced NREM sleep resembles that of spontaneous sleep attacks in *orexin*^-/-^ mice. We found that the activation-induced NREM sleep had an EEG activity comparable to that of spontaneous sleep attacks, further supporting that the activation of SLD^GABA^ neurons induces sleep attacks in *orexin*^-/-^ mice (**Figure S5D-F)**. By comparing the EEG activity, we further validated that activation of SLD^GABA^ neurons did not induce wake→REM sleep transitions, nor did it induce transition into cataplexy in *orexin*^-/-^ mice (**Figure S5A-C and S5G)**. In addition, we found that the activation-induced state could immediately be terminated by tactile stimulation (i.e., touching a mouse with a paintbrush) (**Figure S5H and S5I**). We previously showed an episode of cataplexy cannot be terminated by tactile stimulation^9^. Collectively, our findings demonstrate that activation of SLD^GABA^ neurons induces sleep attacks in *orexin*^-/-^ mice.

Finally, we activated SLD^GABA^ neurons selectively during both NREM and REM sleep in *orexin*^-/-^ mice. Here, we found no difference between genotypes in how activation of SLD^GABA^ neurons changed behaviour during either NREM or REM sleep. As in *orexin*^+/+^ mice, we found that activation of SLD^GABA^ neurons during NREM sleep lengthened NREM sleep and delayed the entrance into wakefulness in *orexin*^-/-^ mice. The magnitude of this effect was similar in both genotypes (**Figure S4A-D)**. Activating SLD^GABA^ neurons during REM sleep had no effect in either *orexin*^+/+^ or *orexin*^-/-^ mice (**Figure S4E and S4F)**. Lastly, we found that neither motor nor cortical activity was impacted by their activation during NREM and REM sleep in either *orexin*^+/+^ or *orexin*^-/-^ mice **(Figure S4G-N)**. These data, therefore, indicate that the activation of SLD^GABA^ neurons reinforces NREM sleep specifically.

### Silencing SLD^GABA^ neurons terminates sleep attacks by promoting wakefulness in *orexin*^-/-^mice

Next, we wanted to determine if silencing SLD^GABA^ neurons can rescue narcoleptic animals from sleep attacks **(Figure 5A)**. First, we investigated whether wakefulness generated by silencing SLD^GABA^ neurons differs between *orexin*^-/-^ and *orexin*^+/+^ mice. We found that silencing SLD^GABA^ neurons had similar wake-promoting effects in both *orexin*^+/+^ and *orexin*^-/-^ mice. As in *orexin*^+/+^ mice, we found that silencing SLD^GABA^ neurons triggered immediate wakefulness from both NREM and REM sleep in *orexin*^-/-^ mice and the latency to wakefulness did not differ between the genotypes (**Figure 5B and 5C**). In addition, we found that the wakefulness triggered by silencing SLD^GABA^ neurons had cortical activity and motor activity that was comparable between *orexin*^-/-^ and *orexin*^+/+^ mice (**Figure S6A-H)**. We also found that silencing SLD^GABA^ neurons in *orexin*^-/-^ mice prolonged wakefulness and increased the probability of wakefulness across time. The magnitude of this effect was not significantly different from that observed in *orexin*^+/+^ mice (**Fig S6I and S6J**). Finally, we found that silencing SLD^GABA^ neurons during wakefulness enhanced both cortical and motor indices of wakefulness to the same extent in both *orexin*^-/-^ and *orexin*^+/+^ mice (**Figure S6K-P**). Together, these data show that the silencing SLD^GABA^ neurons promotes wakefulness and the cortical and motor features that support it, but these effects are unaltered by the loss of orexin signaling.

**Figure 5.**
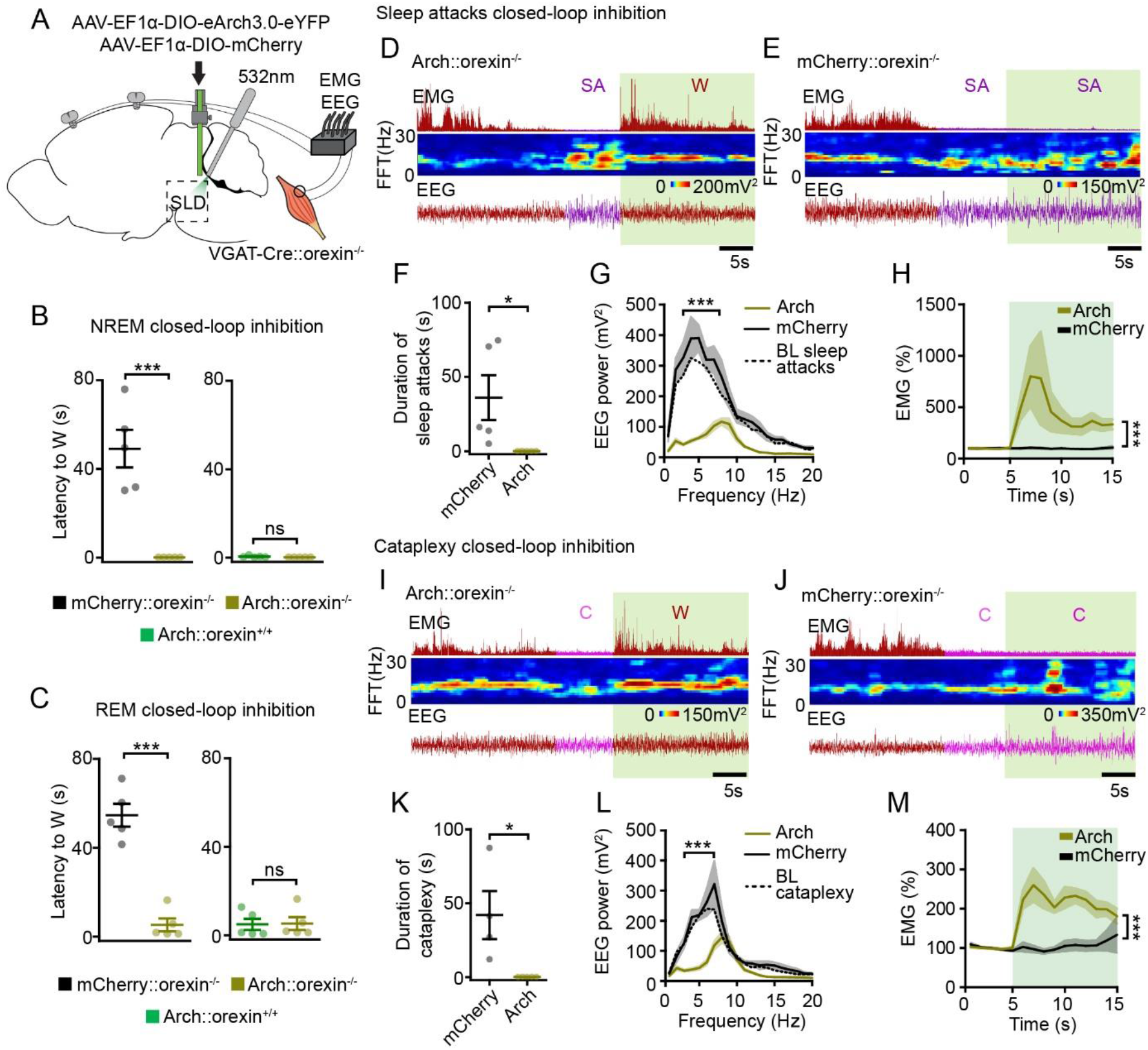
Silencing SLD^GABA^ neurons in orexin^-/-^ mice rescues the animals from sleep attacks and cataplexy. **(A)** A schematic of optogenetic silencing of SLD^GABA^ neurons coupled with EEG and EMG recordings in orexin^-/-^ mice. **(B)** Latency to wake from NREM sleep upon laser stimulation (mCherry::orexin^-/-^ n=5, Arch::orexin^-/-^ n=5, and Arch::orexin^+/+^ n=5; unpaired t-test). **(C)** Latency to wake from REM sleep upon laser stimulation (mCherry::orexin^-/-^ n=5, Arch::orexin^-/-^ n=5, and Arch::orexin^+/+^ n=5; unpaired t-test). **(D-E)** Example polysomnograms with closed-loop stimulation of Arch-expressing and mCherry-expressing SLD^GABA^ neurons during sleep attacks. Shown are EMG amplitude, EEG spectrogram, and EEG raw traces. **(F)** Duration of sleep attacks upon laser stimulation (mCherry n=5 and Arch n=5; unpaired t-test). **(G)** Power spectral density of EEG in response to the laser stimulation during sleep attacks (mCherry n=5 and Arch n=5; Two-way RM ANOVA with Bonferroni post-tests). **(H)** EMG activity before and during laser stimulation from sleep attacks (mCherry n=5 and Arch n=5; Two-way ANOVA with Bonferroni post-test). **(I-J)** Example polysomnograms with closed-loop stimulation of Arch-expressing and mCherry-expressing SLD^GABA^ neurons during cataplexy. **(K)** Duration of cataplexy upon laser stimulation (mCherry n=5 and Arch n=5; unpaired t-test). **(L)** Power spectral density of EEG in response to the laser stimulation during cataplexy (mCherry n=5 and Arch n=5; Two-way RM ANOVA with Bonferroni post-tests). **(M)** EMG activity before and during laser stimulation from cataplexy (mCherry n=5 and Arch n=5; Two-way ANOVA with Bonferroni post-test). Green patches indicate time of laser stimulation. All error bars and shades represent ±s.e.m. *\* p<0.05, ** p<0.01, *** p<0.001 indicate significant differences.*

Because the activation of SLD^GABA^ neurons promotes sleep attacks in narcoleptic mice, we wanted to test whether silencing these neurons could rescue animals from sleep attacks. We tested this by silencing SLD^GABA^ neurons at the onset of individual sleep attacks in *orexin*^-/-^ mice. We found that silencing SLD^GABA^ neurons immediately rescued the animals from sleep attacks into alert and motorically engaged wakefulness, which was not observed in the mCherry control group (**Figure 5D-5H**). These findings, in combination with optical excitation data (**Figure 4**), indicate that SLD^GABA^ neurons are both necessary and sufficient to generate sleep attacks.

Lastly, we wanted to determine if silencing SLD^GABA^ neurons could also rescue narcoleptic animals from cataplexy. Surprisingly, we found that silencing SLD^GABA^ neurons at the onset of cataplexy immediately rescued the animal into wakefulness (**Figure 5I and 5K**) by reinstating brain and motor activity (**Figure 5L and 5M**). Such behavioural effect was not observed in the mCherry control group (**Figure 5J-M**). Because we showed that silencing SLD^GABA^ neurons enhances motor activity in both *orexin*^-/-^ and *orexin*^+/+^ mice (**Figure 1Q & Figure S6P**), we propose that silencing SLD^GABA^ neurons rescues cataplexy by engaging motor activity.

## DISCUSSION

Our study is the first to characterize the functional role of SLD^GABA^ neurons in sleep-wake control and establishes their link to the pathophysiology of narcolepsy. Through precise optogenetic manipulation, we found that SLD^GABA^ neurons function to suppress wakefulness (**Figure 1 & 2**). Next, we found that extensive projections to wake-promoting brain regions, positions SLD^GABA^ neurons as a key nucleus in the neural circuits controlling arousal state transitions (**Figure 3**). Finally, we demonstrated that activation of SLD^GABA^ neurons induces pathological sleepiness, manifesting as sleep attacks in narcoleptic animals and that silencing of these neurons rescue them from this debilitating symptom (**Figures 4, 5 & 6**). While the SLD is recognized as the core center for REM sleep control^8,14,17,39,40^, it has been suggested to be critical for regulating other arousal states. GABA neurons in the SLD display distinct firing patterns across the sleep-wake cycle^21,41^, but their causal role in arousal state control remains speculative. By using cell-type-specific optogenetics in genetically defined populations, we demonstrate that SLD^GABA^ neurons function primarily to suppress wakefulness rather than promote specific sleep states. Several lines of evidence support this conclusion. First, silencing SLD^GABA^ neurons promotes wakefulness from NREM and REM sleep, while activating the same neurons promotes NREM sleep from wakefulness (**Figure 1 & 2**), indicating that their influence extends beyond any single sleep stage. Second, silencing these neurons during existing wakefulness enhances arousal quality by increasing cortical gamma activity and motor engagement (**Figure 1 & 2**). These lines of evidence indicate that the causal role of SLD^GABA^ neurons is to suppress wakefulness and its active characteristics while promoting NREM sleep.

This wake-suppressive function aligns with the broader organizational principles of sleep-wake control, where inhibitory mechanisms balance excitatory arousal systems to enable stable state transitions^14,42–45^. The SLD^GABA^ neurons appear to function as a “brake” on wakefulness, allowing controlled transitions into sleep states when arousal drive diminishes. This regulatory mechanism may be particularly important during vulnerable periods when wake stability is challenged, such as during sleep deprivation, circadian misalignment or during pathologic conditions like narcolepsy^15,46,47^

Our genetically-assisted anatomical mapping reveals that SLD^GABA^ neurons possess the connectivity to modulate multiple aspects of arousal. We show that SLD^GABA^ neurons send axonal projections to various wake-promoting nuclei across the brain^24–29^ (**Figure 3**). These anatomical connections suggest that SLD^GABA^ neurons can simultaneously suppress cortical activation via the medial septum and basal forebrain, and the general wakefulness drive via the lateral hypothalamus and the tuberomammillary nucleus^24,25,27,28,48^. The convergence of these inhibitory inputs likely accounts for the potent wake-suppressive effects we observed. In addition, projections to motor-related areas, including the vestibular nucleus and the spinal cord, may explain their ability to modulate motor activity during wakefulness^31^. These multi-target inputs to both cortical and motor nuclei could ensure stable state control and may prevent partial arousal states that would destabilize sleep-wake boundaries.

Previous work has shown that the SLD also receives inputs from the cortex, forebrain, midbrain, pons and hindbrain^49^ and these inputs may interact with SLD^GABA^ neurons to modulate wakefulness. Finally, the SLD^GABA^ neurons potentially receive inputs from the neighbouring glutamate neurons within the SLD (SLD^GLUT^ neurons). These neurons promote REM sleep and control aspects of the state^9–11,19,41,50,51^. Hence, integration at the level of the SLD may coordinate both REM sleep initiation with wake suppression, ensuring smooth state transitions without arousal interference.

The ability of SLD^GABA^ neurons to suppress wakefulness can be beneficial for sleep stability and normal sleep-wake regulation, but it can also pose a threat to wakefulness stability when engaged at the wrong time as seen in narcoleptic patients^52^. The hypothalamic orexin system normally provides strong excitatory drive to maintain wake stability^33,53^. We found that in orexin-deficient mice (*orexin*^-/-^), activation of SLD^GABA^

neurons triggers rapid sleep attacks—abrupt intrusions of NREM sleep during active wakefulness that closely resemble the clinical presentation of human narcolepsy (**Figure 4**). The activation of SLD^GABA^ neurons in *orexin*^-/-^ mice rapidly initiates NREM sleep even amid active waking behaviour (i.e., eating and moving), bypassing stereotypical sleep-preparatory behaviours (e.g., grooming and nesting) **(Video S2)**. This effect is significantly enhanced compared to wild-type animals, suggesting that orexin loss removes a critical brake on SLD^GABA^ neuron function (**Figure 4**). We hypothesize that when orexin signalling is lost, SLD^GABA^ neurons override residual wake-promoting signals, creating windows of wake instability and triggering sleep intrusions. This hypothesis explains why narcoleptic patients experience sleep attacks even during stimulating activities. The loss of orexin signals allows wake-suppressive mechanisms to dominate inappropriately.

Finally, our discovery that silencing SLD^GABA^ neurons can rescue both sleep attacks and cataplexy (**Figure 5**) opens new therapeutic avenues for narcolepsy treatment. These two cardinal symptoms of narcolepsy—sleep attacks and cataplexy—are distinct physiological states^37^. The ability of SLD^GABA^ silencing to terminate both of them suggests these neurons may serve as a final common neural pathway. While SLD^GLU^ neurons drive cataplexy through REM sleep muscle atonia mechanisms, we propose that the SLD^GABA^ neurons interact with SLD^GLU^ neurons to generate cataplexy. Taken together, we demonstrate that the SLD is the underlying brain structure that causes both cardinal symptoms of narcolepsy (i.e, sleep attacks and cataplexy), making this structure an essential target for future treatment of narcolepsy.

Our study provides definitive evidence for SLD^GABA^ neuron function; however, several important questions remain. First, the precise mechanisms by which orexin loss potentiates SLD^GABA^ function require further investigation. While outside the aim of our current study, direct recordings from SLD^GABA^ neurons in orexin-deficient animals could reveal whether orexin loss alters their intrinsic excitability, synaptic inputs, or both. Second, the upstream signals that normally regulate SLD^GABA^ activity in healthy animals remain unclear. Identifying these regulatory mechanisms could reveal how environmental and circadian factors influence sleep-wake stability. Finally, the thorough investigation of the interaction between the SLD^GABA^ and SLD^GLU^ neurons in both health and disease state would identify what makes the SLD essential for the regulation of the sleep-wake cycle.

## ACKNOWLEDGEMENTS

This work was funded by a grant obtained by J.H.P., the Natural Sciences and Engineering Research Council of Canada (NSERC, 211580). In addition, H.L. received support from the Ontario Graduate Scholarship, Canadian Sleep society, Canadian Institutes of Health Research, and the Canadian Sleep and Circadian Network.

## AUTHOR CONTRIBUTION

conceptualization, H.L., J.J.F., J.H.P. (equal); methodology, H.L., J.J.F. (equal); software, H.L., J.J.F. (equal); formal analysis, H.L.; investigation, HL; resources, J.H.P.; data curation, H.L.; writing – H.L. (lead), J.J.F., J.H.P.; visualization, H.L.; supervision, J.J.F., J.H.P. (lead); project administration, J.H.P.; funding acquisition, J.J.F., J.H.P. (lead)

## DECLARATION OF INTERESTS

The authors declare no competing interests.

## DATA AND MATERIALS AVAILABILITY

All data are available in the main text or the supplementary material

## Notes

### Competing Interest Statement

The authors have declared no competing interest.

